# Zebra finch introductory notes do not influence female response to song and juvenile song learning accuracy

**DOI:** 10.1101/2025.04.06.647492

**Authors:** Shikha Kalra, Chitvan Chandolia, Mahima Gautam, Raghav Rajan

## Abstract

Many songbirds begin their song bouts with introductory notes (INs). In some songbird species like rufous-sided towhees and white-crowned sparrows, INs are believed to function in communication, to alert conspecifics about song onset or as a signal for species identification, but it is unclear whether this function is conserved in other species. Here, we addressed this question using the zebra finch, a well-studied songbird. Male zebra finches begin their song bouts with several INs and the number of INs increases when males sing to females or juveniles, suggesting a possible role of INs in female song preference and/or juvenile song learning. To test these hypotheses, we first recorded and analyzed call responses of female zebra finches to playbacks of different song stimuli. Female zebra finches responded equally to songs with and without INs, even when songs were degraded or when background noise was present. Second, we examined song learning accuracy of juveniles and found that juveniles copied songs accurately from their fathers independent of the fathers’ number of INs. Overall, these results show that zebra finch INs do not influence female response to song or juvenile song learning and suggest that INs might have different functions in different songbird species.

## INTRODUCTION

The song of an adult song bird consists of a stereotyped sequence of sounds (syllables) interleaved by silent gaps and is part of the female-directed courtship ritual of the male [1–5]. The song bouts of many songbird species begin with introductory notes (IN) before the actual song is produced [6–17]. Two broad categories of functions have been hypothesized for INs; (1) communicative [18,19] and (2) motor preparatory functions [20,21].

Communicative functions of INs include their possible function as an alerting signal before the main message [18] or as a signal that guides learning the correct species-specific song [19]. Songs are known to have communicative functions and adult birds often respond to conspecific songs played back through a speaker. Richards [18] tested the responses of Rufous-sided towhees (a songbird species) to modified song playbacks. He found that birds responded strongly to playback of normal songs (without INs) confirming the communicative value of song. However, if these songs were degraded, then towhees only responded if the playback includes normal INs before the degraded songs. This suggested that INs act as an alerting signal before the main message (the song) [18]. Just like adults, juvenile songbirds also respond to playbacks of songs from speakers [22]. If these juveniles have been isolated from other conspecifics early in life, they also learn songs from such playbacks [11,23].

While learning from such playbacks, in the absence of a live bird, juveniles need to decide whether the played back songs are songs of their own species or not. Soha and Marler showed that juvenile white-crowned sparrows decide whether to learn from a playback or not based on the presence of an introductory whistle that is characteristic of normal white-crowned sparrow songs [9,19]. Juveniles will even learn the songs of other species, if they are preceded by the normal white-crowned sparrow introductory whistle [19]. These two examples highlight possible communicative functions of INs.

In the zebra finch, a well-studied songbird species, INs are hypothesised to perform a motor preparatory function [20,21]. Male zebra finches begin their song bouts with a variable number of INs before producing multiple renditions of their songs. Across song bouts, the timing between successive INs becomes faster and more stereotyped as INs get closer to the last IN, just before song. Successive INs also become louder and more consistent in their acoustic properties. Along with changes in timing and acoustic properties, premotor neural activity during INs, also reaches consistent firing patterns just before the last IN. Thus behaviourally and neurally, INs progress from a variable “initial” state during the first IN to a consistent “final” state during the last IN, just before each song begins [20]. This progression is independent of two forms of sensory feedback [21]. Given the similarities of this IN progression to the progression of neural preparatory activity in primates and rodents, from a variable initial state to a more consistent state just before movement initiation, INs have been suggested to reflect the songbird brain “preparing” to produce song.

While INs have been hypothesized to function as motor preparation in the zebra finch, previous studies have shown that male zebra finches produce more INs before their songs when singing to a female [24] or a juvenile [25], as compared to when they sing alone.

During female-directed courtship, male zebra finches produce more INs at the start of their song bouts [24] (Fig. 1A) and in song preference assays, female birds prefer female-directed courtship songs over “alone” songs (songs produced in isolation). This is apparent in the increased time spent near speakers that produce female-directed courtship songs [26]. The increase in mean number of INs in the courtship context is not learned and is present in birds raised without exposure to other adult males (isolates; [27]). Similar to female-directed courtship songs, males also sing more INs at the start of song bouts directed towards juvenile birds [25] (Fig. 1A). Juveniles that pay more “attention” to tutor songs show greater learning as seen by their increased similarity to tutor songs [25]. These results suggest that the increase in number of INs before song in both these contexts could serve to increase the attention of the receiver to the song that follows. This suggests that zebra finch INs serve a communicative function in both these contexts similar to the hypothesized communicative function of INs in rufous-sided towhees [18] and white-crowned sparrows [19].

**FIGURE 1.**
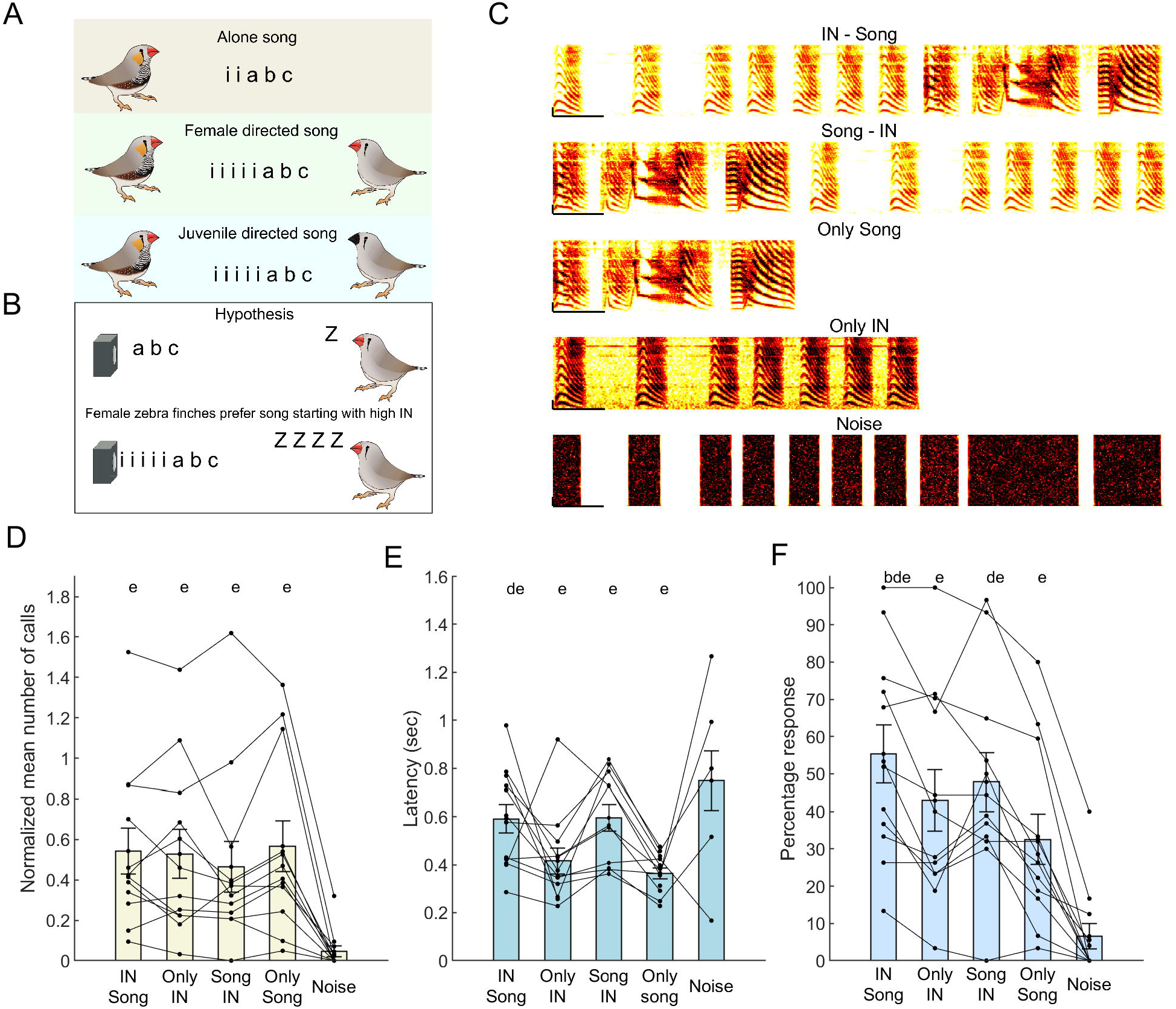
Female zebra finches respond with equal number of calls to playback of songs with and without INs before them. (A) Cartoon summarizing previous research that shows that male zebra finches sing more introductory notes (represented by ‘i’) when they sing to a female (female-directed song) and when they sing to a juvenile (juvenile-directed song), compared to songs produced when the bird is alone. (B) Cartoon summarizing our hypothesis that female zebra finches produce more calls in response to songs preceded by INs as compared to songs without any introductory notes (‘i’ represents introductory notes, ‘a’, ‘b’, and ‘c’ represent song syllables and ‘z’ represents female calls). (C) Spectrogram showing the 5 different stimuli that were played back. Scale bar - x-axis - 150ms; y-axis - 1 kHz. (D), (E) and (F) Response of female zebra finches to the different playback stimuli. Mean number of calls normalized to stimulus duration (D), latency of response (E) and percentage of trials with a response (F) are plotted for each stimulus. Each circle represents one bird and lines connect data from the same bird. Bars represent mean and s.e.m across all birds (n=12). ‘e’ represents significant difference from noise, ‘b’ represents significant difference from only IN stimulus and ‘d’ represents significant difference from only song stimulus, p < 0.05, Repeated measures ANOVA followed by post-hoc Tukey Kramer test. Note: for the latency (E), there are no points for birds that did not call during any of the trials of a particular type.

Here, we tested whether INs have a communicative role in the context of female-directed and juvenile-directed songs. First, we played back songs, with and without INs, to female zebra finches and analyzed their responses. We hypothesized that females would respond more to songs preceded by INs (Fig.1B). Second, we examined whether the accuracy of song learning in juveniles was correlated with the mean number of INs produced by fathers, at the start of their song bouts.

## MATERIALS AND METHODS

All procedures at IISER Pune were conducted after approval from the Institute Animal Ethics Committee (IAEC), IISER Pune, and were in accordance with guidelines of CCSEA (Committee for the Control and Supervision of Experiments on Animals), New Delhi, India.

### Experimental birds

Adult female zebra finches (>90 days post-hatch; n=27 birds, 24 lab bred and 3 bought from outside vendor) were used for playback experiments. 8 of these birds were excluded from the analyses as they did not respond for more than 1 day. 65 male birds and their fathers (16 fathers) from 16 nests were used for testing song learning.

### Playback and experimental design

Before initiating the playback experiment, female zebra finches were isolated for 2 days in a sound attenuation chamber in separate cages (2 birds per box separated by cardboard partition). Playback experiments were done in a cage (dimension: 38 x 27 x 29 cm), placed in a sound-attenuation enclosure (NewTech Engineering Systems, Bangalore, India) with the playback speaker in front of the cage, through which all stimuli were played. For the experiments with noise playback, an additional speaker was placed on top of the cage and noise was played back through both this additional speaker and the playback speaker. On the day of the experiment, birds were kept in the setup for acclimatization, 15-30 minutes prior to starting stimulus playback. Following this acclimatization, playback started using a custom MATLAB script. We conducted 1 session per day, where each session consisted of two trials separated by 10 minutes (for the first set of 4 birds, we conducted 3 sessions per day separated by 6 minutes and then changed this as there seemed to be fewer responses for the 2nd and 3rd session). For all experiments, each trial consisted of the different stimulus types played in random order with an inter-stimulus gap of 90 seconds and each stimulus type consisted of 3 repeats of the stimulus with 1.5 seconds silence in-between successive repeats. For the second set of experiments with background noise, we randomly chose one of the trials in the session to play background noise; the other trial was without background noise. Background noise began 10 minutes before the first stimulus and ended after the last stimulus type was played. We recorded sounds from our zebra finch colony and played back these sounds as background noise to mimic a noisy zebra finch colony environment. We used a decibel meter to measure the sound pressure levels for the noise and the stimulus. Over a 60 second period, the maximum sound pressure level at the centre of the cage was 76.4 dB when we played the song and IN stimulus and the background noise had a maximum sound pressure level of 68.4 dB, keeping the stimulus just audible over the background. Noise was played back through both the speakers (to provide an illusion of noise from everywhere) whereas only the second speaker was used to playback stimuli (to provide an illusion of a localized playback).

For a given bird, playback experiments were conducted for 3-5 days (5 days for experiment 1 and 3-4 days for the background noise experiment), interleaved by one day, to prevent habituation (different birds were used for the two experiments). Calls produced by the female and songs played back were recorded by placing an omnidirectional microphone (AKG C417PP) on top of the bird’s cage, towards the front. Signals from the microphone were amplified, band-pass filtered (Behringer Xenyx 802), digitized, and recorded on a computer at a sampling rate of 44100 Hz using a custom Python program. We also used a webcam to record videos of the behaviour of the female (30 frames per second, 1280 x 720 resolution) during the experiment with background noise. The webcam was placed above and behind the playback speaker in front of the cage. For these experiments, the timing of playback stimuli were determined from the webcam recording as the microphone recording was not very clear due to the background noise.

### Stimuli used for playback

Songs from one adult male zebra finch (randomly chosen from our colony) was used to generate different stimuli for the playback experiments. For the first experiment, we used five different stimuli (Fig. 1C) namely, (1) normal zebra finch song (7 INs followed by a song),(1) only the song, (3) only 7 INs, (4) reverse song (song followed by 7 INs) and, (5) white noise. The white noise stimulus had the same temporal pattern as the normal song (7 INs and motif) but the acoustic features were replaced with white-noise generated using the MATLAB (MathWorks) function wgn (White Gaussian noise). Reverse song and noise were used as controls; reverse song was a control for the duration of normal INs + normal song, while noise was used as a negative control. For the background noise experiment, the song of a bird from another colony (recorded at UCSF) was used to make different stimuli. A total of five stimuli were used (Fig. 2A) namely: 1) Normal IN – normal song, 2) Only normal song, 3) Degraded IN – degraded song, 4) Normal IN – degraded song and 5) Only degraded song. Each stimulus consisted of two repeats of the song. The degraded part of the song was generated by randomly shuffling the frequencies of the normal recorded song. To do so, MATLAB function stft (short-time fourier transform) was used to calculate frequencies of the signal at each time point (which depends upon the sampling rate of the song). Frequencies were randomly shuffled for a given time point to preserve the amplitude envelope of the original song. MATLAB function istft (inverse short-time fourier transform) was then used to convert the frequency data back to a time-varying signal. This signal was converted to an audio playback using MATLAB function audiowrite.

**FIGURE 2.**
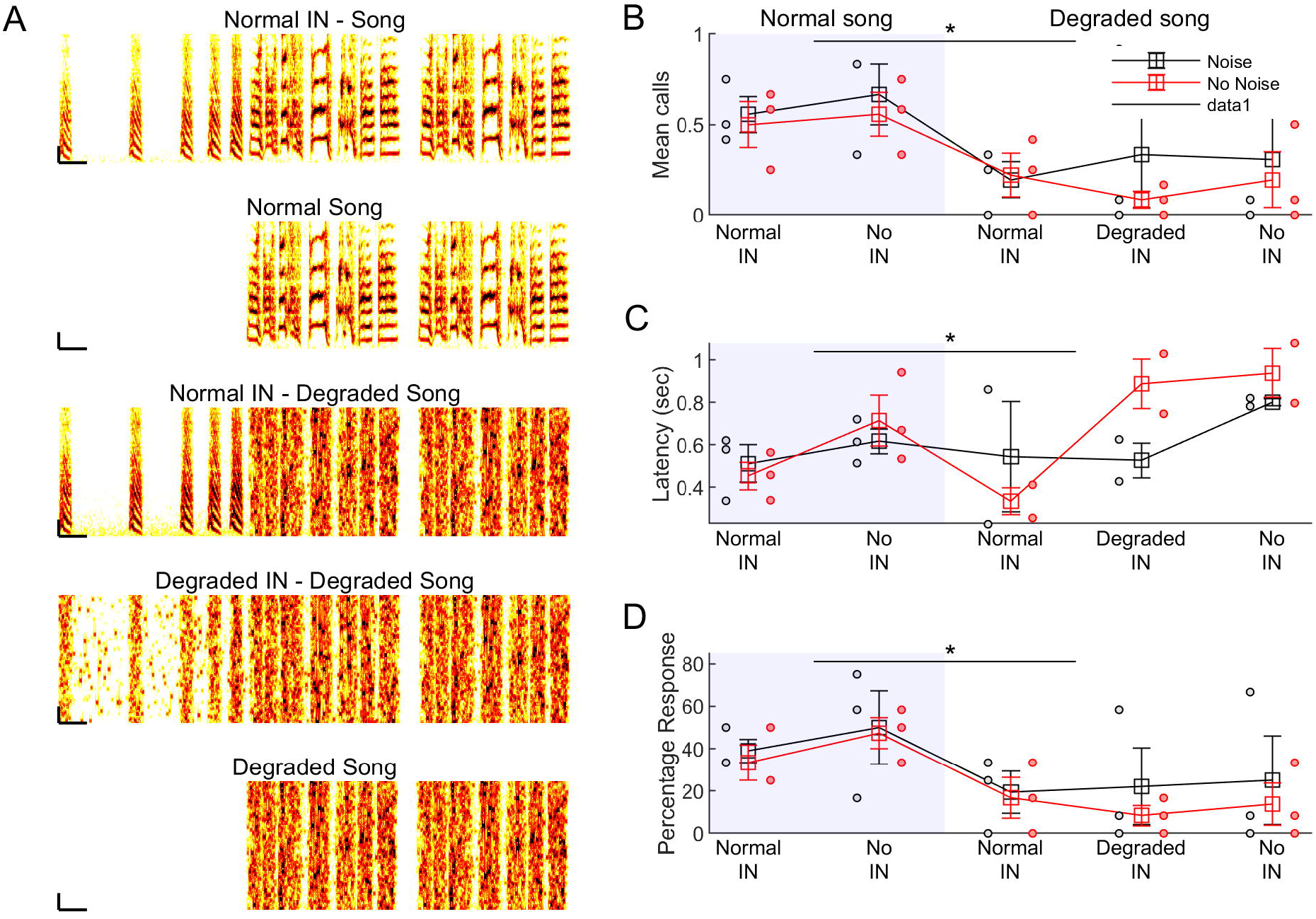
Female zebra finches respond more to normal songs as compared to degraded songs independent of IN number and background noise levels. (A) Spectrogram showing the 5 different stimuli that were played back in the absence of noise. The same stimuli were also played back in the presence of noise. Scale bar - x-axis - 150ms; y-axis - 1 kHz. (B), (C) and (D) Response of female zebra finches during the song part of the different playback stimuli. Mean number of calls produced only during the song (B), latency of response (C) and percentage of trials with a response (D) are plotted for each stimulus in the different conditions. Shaded region represents normal song, while region without shading represents degraded song. Black lines represent presence of background noise and red lines represent absence of background noise. Squares and whiskers represent mean and s.e.m across all birds (n=3). Black and red circles represent data from individual birds (red for trials with no background noise and black for trials with background noise). * represents significant difference between normal song and degraded song (p<0.05 for Song quality, Linear Mixed Effects Model)

### Analysis

All analysis was done using custom-written scripts in MATLAB (MathWorks, panel function was used for plotting [28]). Playback syllables and calls produced by the females were manually segmented and annotated using custom written MATLAB scripts. For playback for experiment 1, we annotated the first syllable of the stimulus (IN or song syllable depending on the stimulus being played back) to mark the onset and category of the stimulus played back. For experiment 2 with background noise, it was difficult to identify the first syllable within the background noise and so we used the first or last IN for the normal INs + normal song stimulus, last syllable offset for the degraded INs + degraded song stimulus, first syllable for the normal song stimulus, last syllable offset for degraded song stimulus and third IN or last IN for normal INs + degraded song stimulus. This was used as a marker and the time of stimulus onset was calculated based on this for each stimulus type. The number of calls produced by the female during the stimulus period was counted as a measure of the response. The number of calls produced over each repetition of the stimulus were normalized to the duration of the stimulus and then averaged across all repetitions across all of the playback days. Response latency was calculated as the interval between the onset of the stimuli and the onset of the first call produced by the female zebra finch. For birds that did not call for any of the trials for a particular stimulus type, we used the maximum duration of the stimulus as the latency for the repeated measures ANOVA statistical test. Response percentage was defined as the percentage of trials with atleast one female call. Birds that did not call for more than one session were excluded from the analysis and in some cases, we did not continue more sessions with such birds.

### Comparisons between father and son

Data for the mean number of INs and mean song similarity for fathers and their sons was collected as part of an older study (n=65 birds from 16 nests; [29]). Here, we only used this data to compare the mean number of INs produced by fathers to the accuracy of song copying as measured by the song similarity between son and father [29].

### Statistics

Statistics was done using MATLAB and Python package statsmodels. Repeated measures ANOVA (MATLAB function ranova) was used for the comparison across groups with a significance threshold of 0.05. When the ANOVA comparison was significant, a Tukey-Kramer post-hoc test was used to compare between groups (MATLAB function multcompare). For stimuli, where the birds did not respond on any of the trials, the mean response latency was considered to be the duration of the longest stimulus (maximum latency). For correlation of song copying by juveniles and the mean IN number of the father, Pearson correlation coefficient was calculated (MATLAB function corrcoef). For the background noise experiments, we used a linear mixed effects model (python package statsmodels) with number of calls, or response latency, or percentage of trials with a response, as the response variable and song quality (degraded or normal), IN quality (degraded, normal or absent) and background noise (present or absent) as predictor variables (separate tests were run for each response variable). All statistics were run on the group data with average values representing each bird.

## RESULTS

### The presence of INs before male zebra finch song does not influence females’ preference

To determine the influence of INs on female song preference, we used a call-response assay (n=12 female zebra finches). In this assay, different song stimuli were played back through a speaker to a female zebra finch and the total number of calls produced by the female during the stimulus was recorded as a measure of preference. Calls produced are known to be a reliable measure of preference [30,31]. In addition, in female birds implanted with estradiol, the number of calls produced are correlated with the number of courtship solicitation displays (CSDs) produced, further validating calls as a good measure of song preference [32,33].

Here, we used 5 different stimuli, namely, (1) a normal zebra finch song preceded by 7 INs, (2) a song without INs, (3) 7 INs alone without the song, (4) reverse song, i.e. the song followed by 7 INs, and (5) a noise stimulus where the temporal structure of a normal INs followed by song was maintained, while the spectral structure was changed into white noise (see Methods for details). Across the four different song stimuli, the average number of calls normalised to the duration (Fig. 1D), the response latency (Fig. 1E) and the fraction of trials with a response (Fig. 1F) were similar with only minor differences between specific subsets. When compared to the four song stimuli, noise received fewer calls, longer response latencies and a smaller fraction of trials with responses (Fig. 1D,1E, 1F, n=12, p < 0.05, Repeated measures ANOVA, post-hoc Tukey Kramer Test, Table S1)

The song used in this assay was a song that the females were not familiar with and previous studies have shown that females respond more to familiar songs, like those of their father or their mate [34,35]. To test whether the preferences were influenced by song familiarity, we repeated these experiments with familiar songs; i.e. the song of a father was used to test for preference with 4 female offspring. Again, females produced equal number of calls for songs with and without INs (Fig. S1 n=4 female zebra finches). Overall, these results demonstrate an equal preference for songs with and without INs, suggesting that under normal conditions, INs do not play a role in female song preference.

### INs do not increase female response to degraded songs, even in the presence of background noise

Previous studies in Anoles lizards suggest that introductory gestures might function to attract the attention of a receiver [36]. Anoles lizards extend their dewlap (skin under the throat) as part of their threat display and under noisy visual conditions, i.e. when light levels are low or when there is more visual noise from plants blowing in the wind, they add more introductory push-ups, before the dewlap extension. In response to a robotic lizard performing the threat display, other lizards responded more to threat displays that were preceded by push-ups when the visual environment was noisy [36]. Similar to this result, playback experiments with rufous-sided towhees (a species of songbird) showed that towhees responded more to degraded songs, if they are preceded by INs [18]. These studies suggest that introductory gestures might serve to attract the attention of a receiver, especially under conditions where the main message is harder to detect.

To test if zebra finch INs had a similar function, we repeated the song playbacks with a different set of stimuli (Fig. 2A), in the presence and absence of background noise (n=3 female zebra finches, 3 additional birds were tested but did not call and were excluded from the analysis). Here, we used the sounds recorded in our bird colony to simulate background noise as zebra finches typically live in groups. In addition to the normal song with and without INs, we also included 3 more stimuli where the song was degraded. This was motivated by previous results showing that rufous-sided towhees respond to degraded songs, if they are preceded by INs [18]. Specifically, the 3 additional stimuli were (1) normal INs followed by degraded song, (2) degraded INs followed by degraded song and, (3) degraded song without INs before it. We quantified calls produced only during the song as a measure of female response. Females produced more calls in response to normal songs as compared to degraded songs (Fig. 2B, Table S1, p = 0 for Song quality, Linear Mixed Effects Model). The latency to the first call was shorter for normal songs as compared to degraded songs (Fig. 2C, Table S1, p=0.014 for Song Quality, Linear Mixed Effects Model) and the percentage of trials with a response was also higher for normal songs as compared to degraded songs (Fig. 2D, Table S1, p = 0 for Song quality, Linear Mixed Effects Model). The presence/absence of INs or the presence/absence of background noise did not affect female responses (Fig. 2B, 2C, 2D, Table S1). Overall, these results show that the presence of INs before the song does not increase the response to song, even when the song is degraded or when there is background noise present. These results suggesting that zebra finch INs do not serve as an alerting signal in the context of female-directed courtship song.

### The presence of introductory notes does not influence the accuracy of motif copying

As mentioned earlier, male zebra finches also increase the number of INs when singing to a juvenile bird [25]. We next examined if increased IN number is correlated with the accuracy of song copying by juvenile finches. We found no correlation between the mean number of INs produced by a father and the accuracy of song copying by a son (measured as the similarity between the song of the son and the song of the father) (Figure 3A, n = 65, r = 0.08, p= 0.524, Pearson correlation coefficient). Infact, sons even copied songs accurately from the rare zebra finch fathers without any INs before their songs and the copying accuracy was not different from the accuracy of copying songs from fathers with INs before their songs (Fig. S3, p=0.94, Wilcoxon ranksum test). These results suggests that the presence of INs does not affect the accuracy of song copying by juvenile zebra finches.

**FIGURE 3.**
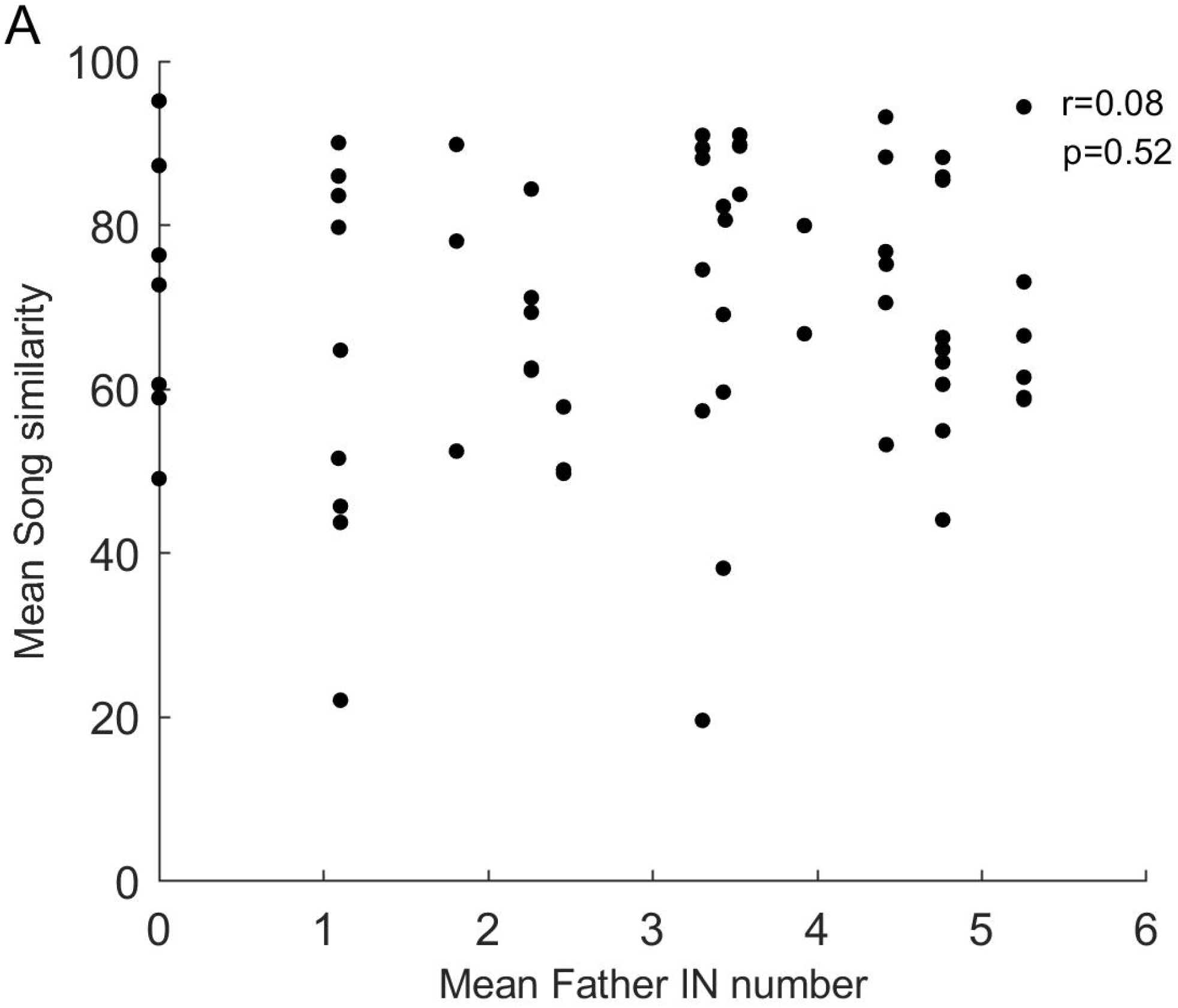
Accuracy to song of the father is not correlated with mean number of INs produced by the father. (A) Mean similarity to song of the father for juveniles (n=65 birds from 16 nests). Each circle represents one bird.

## DISCUSSION

Song bouts of many different songbird species begin with introductory notes (INs). Communicative or motor preparatory functions have been hypothesized for these INs, albeit separately in different songbird species [18–21]. In the zebra finch, a well-studied songbird, a motor preparatory function has been hypothesized for INs [20,21]. However, male zebra finches increase the number of INs when singing to a female or when singing to a juvenile, suggesting a communicative function for INs in these two contexts. Here, we addressed this question using song playback assays with the female zebra finch and by analyzing the accuracy of song copying in juveniles. First, using a song playback assay we showed that female zebra finches produce equal number of calls in response to songs with and without INs (Fig. 1). While females called less to degraded songs, their response was not influenced by the presence of INs, even if there was a noisy background (Fig. 2). Second, we showed that the song copying accuracy of juvenile birds was not affected by the number of INs (Fig. 3). Juveniles also copied songs accurately when tutored by zebra finches that did not produce INs before their songs. These two results demonstrate that zebra finch INs do not function as a communicative signal in the context of female-directed and juvenile-directed songs.

### Function of INs in female-directed courtship songs

Our results show that female zebra finches responded equally to songs with and without INs, suggesting the females do not show greater preferences to songs with INs (Fig. 1 and Fig. 2). Why do males increase the number of INs before female-directed songs? One possible explanation involves INs functioning as motor preparation. In addition to having more INs at the beginning, female-directed songs are also faster and less variable [24,37]. If INs reflect motor preparation for song, it is possible that more INs before female-directed song reflects more preparation for producing faster and less variable songs. Given the preference of female zebra finches for female-directed songs [26], more preparation could result in songs that are more preferred by the female. Future analyses examining the correlations between the number of INs before song and song stereotypy could help test this hypothesis. Alternatively, negative reinforcement paradigms [38] could be used to train birds to increase the number of INs; if birds can increase the number of INs, the song that follows should also be less variable.

Another possible reason for more INs before female-directed songs could be to delay the decision to sing. Producing song involves a little more oxygen consumption than most other activities [39–42] and male birds might decide to sing only if the female is responsive. INs are short duration syllables that are also lower in amplitude and the cost of producing a few INs would be less than the cost of producing a full song. Experiments with strobe light perturbations have shown that birds can only interrupt their songs at the boundaries of syllables [43]. Given that INs are typically shorter than song syllables [11,29], INs might allow birds to quickly stop singing if the female is not responsive. Previous studies have shown that birds sing more motifs if the female responds [44,45], but whether the IN to song transition is dependent on female response is not clear. A more careful examination of the dynamics of courtship song performance along with female response would help to determine if the IN to song transition depends on female response. Alternatively, one could allow male zebra finches to interact with virtual female birds [46–48] and control the response of the virtual female bird to see if progression to song depends on a response from the female.

### Function of INs in juvenile-directed song

Our results suggest that the number of INs sung by a male do not play a role in attracting the attention of the juvenile. While our experiments did not control for variation in songs between fathers, the fact that differences in IN number did not influence copying suggests that INs do not play a role in helping the juvenile pay attention and copy song accurately.

Artificial tutoring experiments [49] with the same song, with and without INs, could confirm that INs do not have any role to play in influencing the accuracy of song copying. A recent study using such artificial tutoring experiments has shown that the mean number of INs before a song bout and IN acoustic properties are learned by young zebra finches from a tutor [29]. This suggests that juveniles treat INs as another syllable and learn the statistics of the entire bout.

### INs could play a role in male-male competition

In frogs, INs have been suggested to function in male–male competition [50]. It is possible that INs produced by male zebra finches also have similar functions. If it is true, one would expect, male zebra finches to produce more INs in the presence of another male and one would expect other adult male zebra finches to respond more strongly to playbacks with greater number of INs. It would be interesting to see if the number of males in the vicinity also influences the number of INs produced. Currently it is difficult to record the vocalizations for multiple individuals without interference. Recent studies have used head implanted microphones [45] or wireless mics [51] to record interacting birds and such an approach could be taken to check if INs have a role in male-male interactions.

Our results rule out two important hypotheses related to the function of INs in the zebra finch and suggest that INs could have other communicative / non-communicative functions. Given that our results are different from the proposed function of INs in other songbird species [18,19], a systematic study of the presence/absence of INs in other songbird species would help understand whether INs are a common feature of the song bouts of all songbirds. The presence or absence of INs could then be correlated with habitat and life history traits of the specific species to make better hypotheses about the function of INs. Overall, our results suggest that INs in the zebra finch do not have a communicative function in the context of female-directed and juvenile-directed songs and suggest the possibility that INs have different functions in different songbird species.

## ACKNOWLEDGMENTS

We would like to thank Prakash Raut for help with bird colony maintenance. We would like to thank members of the Rajan Lab for useful discussions and comments on the manuscript.

## SUPPLEMENTARY INFORMATION

Supplementary information includes 3 supplemental figures and 1 supplemental table.

