## Supplementary material for "Zebra finch introductory notes do not influence female response to song and juvenile song learning accuracy": Fig. S1, Fig. S2, Fig. S3, Table S1

### SUPPLEMENTAL INFORMATION

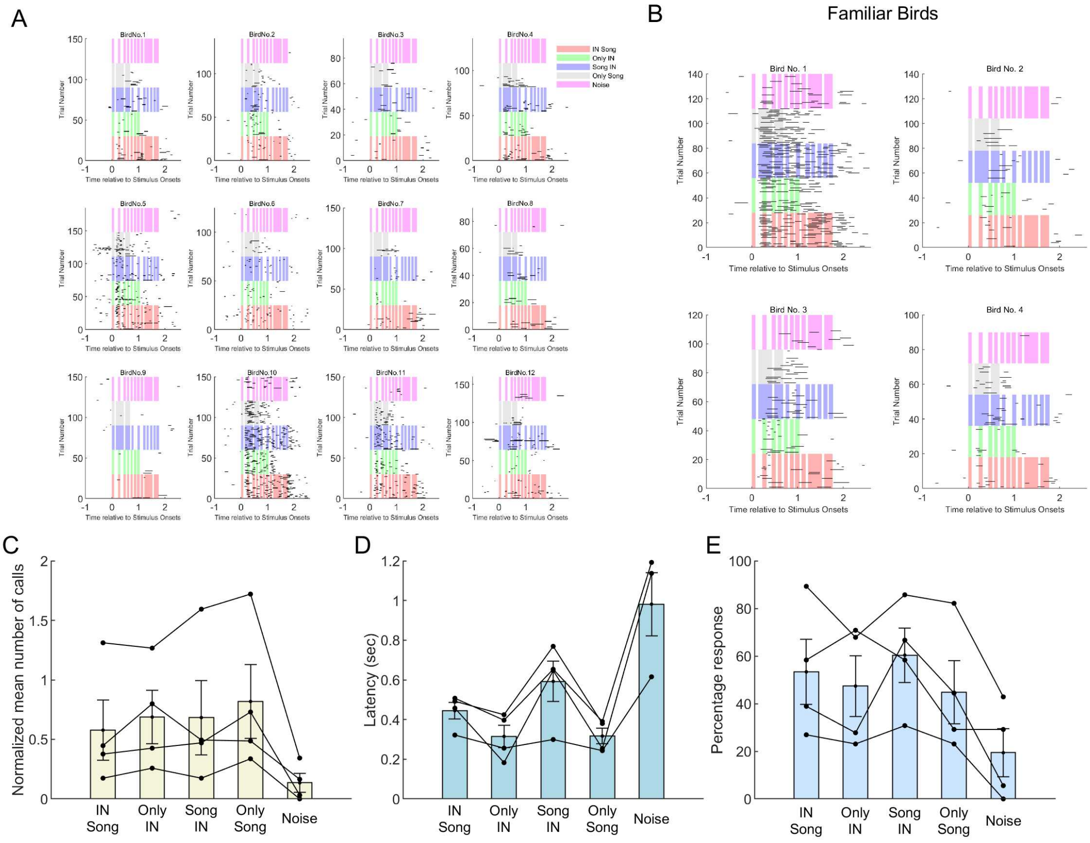


**FIGURE S1 <u>Raster of responses and responses of female birds familiar with the song</u>**

(A) Raster plots showing the time of occurrence of calls during the different stimuli. The colour shading represents different stimuli; orange for INs + song, green for only INs, blue for song + INs, gray for only song and pink for noise. 0 on the x-axis represents time of stimulus onset and each row represents one trial. Black ticks represent time and duration of calls produced by the female. Each plot is for one individual female bird (total n=12 females used for analysis shown in Fig. 1D, 1E and 1F) (B) Raster plots showing the time of occurrence of calls for female zebra finches that were familiar with the song (n=4, song of their father). Labelling and colours are same as (A). Stimuli used were same as the ones shown in Fig. 1C. (C), (D), and (E) Response of female zebra finches to the different playback stimuli for the familiar songs experiment. Mean number of calls normalized to stimulus duration (D), latency of response (E) and percentage of trials with a response (F) are plotted for each stimulus. Each circle represents one bird and lines connect data from the same bird. Bars represent mean and s.e.m across all birds (n=12).

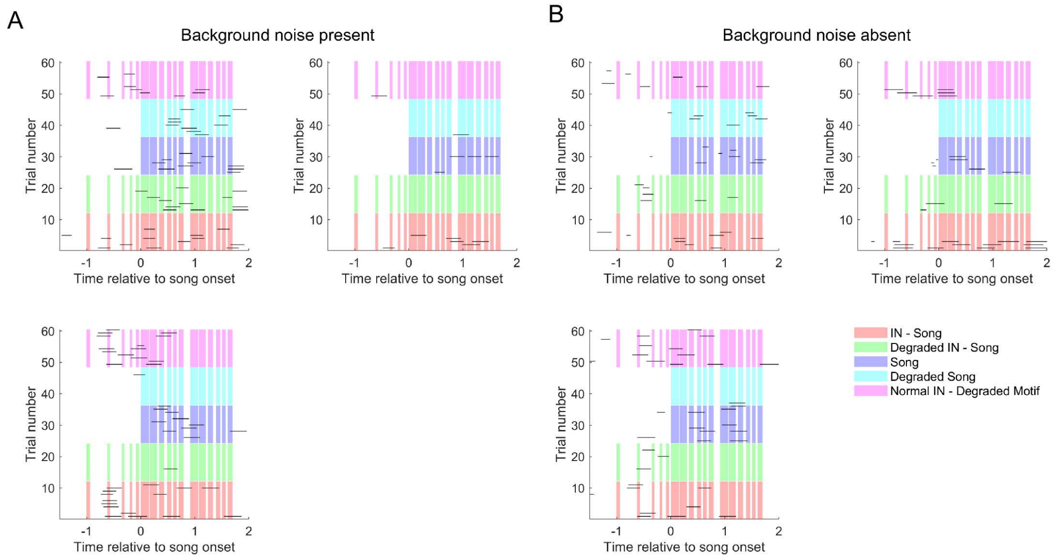


**FIGURE S2 <u>Rasters of responses of female birds to normal/degraded songs with and without background noise.</u>**

(A) and (B) Raster plots showing the time of occurrence of calls during the different stimuli in presence of noise (A) and in the absence of noise (B) for the same set of birds. The colour shading represents different stimuli; orange for INs + song, green for degraded INs + degraded song, blue for normal song alone, cyan for degraded song alone and pink for normal INs + degraded song. 0 on the x-axis represents time of song onset and each row represents one trial. Black ticks represent time and duration of calls produced by the female. Each plot is for one individual female bird (total n=3 females used for the analysis shown in Fig. 2B, 2C and 2D).

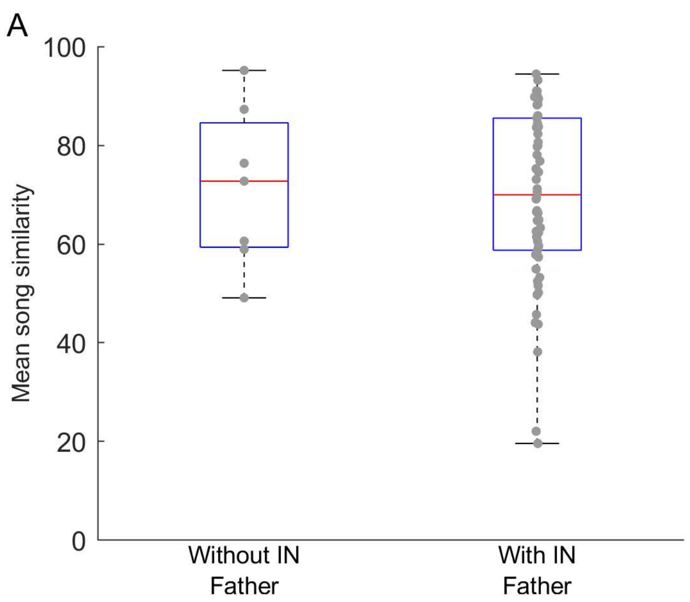


**FIGURE S3 Song similarity to father is similar for juveniles with fathers with INs and without INs**

(A) Box plot showing the song similarity to the father's song for juveniles with fathers who did not sing INs before song and juveniles with fathers who sang INs before their song. Each circle is one juvenile. Same data as in Fig. 3A. $p = 0.94$, Wilcoxon ranksum test.

**TABLE S1 <u>Results of the statistical tests</u>**

Significant differences are highlighted in yellow.

**<u>Statistics for Figure 1</u>**

Number of birds = 12; Number of stimuli = 5

Stimuli order: 1- IN motif; 2- Only IN; 3- Motif IN; 4- Only motif; 5- Noise

A) Normalized mean number of calls (Test used: repeated measures ANOVA (ranova); Posthoc: Tukey Karmer)

| | SumSq | DF | MeanSq | F | pValue | pValueGG | pValueHF | pValueLB |
|---|---|---|---|---|---|---|---|---|
| (Intercept):AllStimuli | 0.38182 | 4 | 0.095456 | 2.7224 | 0.042779 | 0.095155 | 0.081719 | 0.12996 |
| Birdno:AllStimuli | 0.15393 | 4 | 0.038483 | 1.0975 | 0.37101 | 0.35002 | 0.35723 | 0.31947 |
| Error(AllStimuli) | 1.4025 | 40 | 0.035063 | | | | | |

**<u>Posthoc test</u>**

| AllStimuli_1 | AllStimuli_2 | Difference | StdErr | pValue | Lower | Upper |
|---|---|---|---|---|---|---|
| 1 | 2 | 0.014722 | 0.04596 | 0.99732 | -0.13654 | 0.16598 |
| 1 | 3 | 0.078302 | 0.039721 | 0.34397 | -0.052424 | 0.20903 |
| 1 | 4 | -0.024213 | 0.059923 | 0.99347 | -0.22142 | 0.173 |
| 1 | 5 | 0.49777 | 0.096256 | 0.0029772 | 0.18099 | 0.81456 |
| 2 | 1 | -0.014722 | 0.04596 | 0.99732 | -0.16598 | 0.13654 |
| 2 | 3 | 0.06358 | 0.050105 | 0.71446 | -0.10132 | 0.22848 |
| 2 | 4 | -0.038935 | 0.03682 | 0.82347 | -0.16011 | 0.082244 |
| 2 | 5 | 0.48305 | 0.11028 | 0.0093979 | 0.12012 | 0.84598 |
| 3 | 1 | -0.078302 | 0.039721 | 0.34397 | -0.20903 | 0.052424 |
| 3 | 2 | -0.06358 | 0.050105 | 0.71446 | -0.22848 | 0.10132 |
| 3 | 4 | -0.10252 | 0.060027 | 0.47094 | -0.30007 | 0.09504 |
| 3 | 5 | 0.41947 | 0.09901 | 0.011663 | 0.093618 | 0.74532 |

| | | | | | | |
|---|---|---|---|---|---|---|
| 4 | 1 | 0.024213 | 0.059923 | 0.99347 | -0.173 | 0.22142 |
| 4 | 2 | 0.038935 | 0.03682 | 0.82347 | -0.082244 | 0.16011 |
| 4 | 3 | 0.10252 | 0.060027 | 0.47094 | -0.09504 | 0.30007 |
| 4 | 5 | 0.52198 | 0.11162 | 0.0060618 | 0.15463 | 0.88934 |
| 5 | 1 | -0.49777 | 0.096256 | 0.0029772 | -0.81456 | -0.18099 |
| 5 | 2 | -0.48305 | 0.11028 | 0.0093979 | -0.84598 | -0.12012 |
| 5 | 3 | -0.41947 | 0.09901 | 0.011663 | -0.74532 | -0.093618 |
| 5 | 4 | -0.52198 | 0.11162 | 0.0060618 | -0.88934 | -0.15463 |

==========================================================

B) Latency

Calls produced only during stimulus are considered. In case birds didn't respond during any of the trials for a particular stimulus type, mean latency of 1.77 sec was used for statistics, since that was the maximum possible latency.

| | SumSq | DF | MeanSq | F | pValue | pValueGG | pValueHF | pValueLB |
|---|---|---|---|---|---|---|---|---|
| (Intercept):AllStimuli | 4.4397 | 4 | 1.1099 | 14.471 | 2.1638e-07 | 0.00010375 | 1.7693e-05 | 0.0034629 |
| Birdno:AllStimuli | 1.1312 | 4 | 0.28279 | 3.6869 | 0.012001 | 0.041415 | 0.028956 | 0.08379 |
| Error(AllStimuli) | 3.068 | 40 | 0.076701 | | | | | |

**<u>Posthoc test</u>**

| AllStimuli_1 | AllStimuli_2 | Difference | StdErr | pValue | Lower | Upper |
|---|---|---|---|---|---|---|
| 1 | 2 | 0.17534 | 0.084149 | 0.29746 | -0.1016 | 0.45228 |
| 1 | 3 | -0.10189 | 0.081729 | 0.72664 | -0.37086 | 0.16709 |
| 1 | 4 | 0.22721 | 0.054249 | 0.012546 | 0.048676 | 0.40575 |
| 1 | 5 | -0.75287 | 0.12908 | 0.0012057 | -1.1777 | -0.32806 |
| 2 | 1 | -0.17534 | 0.084149 | 0.29746 | -0.45228 | 0.1016 |
| 2 | 3 | -0.27723 | 0.12532 | 0.25028 | -0.68967 | 0.13522 |

| | | | | | | |
|---|---|---|---|---|---|---|
| 2 | 4 | 0.051872 | 0.048593 | 0.81871 | -0.10805 | 0.21179 |
| 2 | 5 | -0.92822 | 0.15799 | 0.0011396 | -1.4482 | -0.40826 |
| 3 | 1 | 0.10189 | 0.081729 | 0.72664 | -0.16709 | 0.37086 |
| 3 | 2 | 0.27723 | 0.12532 | 0.25028 | -0.13522 | 0.68967 |
| 3 | 4 | 0.3291 | 0.1142 | 0.093981 | -0.046725 | 0.70493 |
| 3 | 5 | -0.65099 | 0.12496 | 0.0028219 | -1.0622 | -0.23973 |
| 4 | 1 | -0.22721 | 0.054249 | 0.012546 | -0.40575 | -0.048676 |
| 4 | 2 | -0.051872 | 0.048593 | 0.81871 | -0.21179 | 0.10805 |
| 4 | 3 | -0.3291 | 0.1142 | 0.093981 | -0.70493 | 0.046725 |
| 4 | 5 | -0.98009 | 0.15095 | 0.00051565 | -1.4769 | -0.48331 |
| 5 | 1 | 0.75287 | 0.12908 | 0.0012057 | 0.32806 | 1.1777 |
| 5 | 2 | 0.92822 | 0.15799 | 0.0011396 | 0.40826 | 1.4482 |
| 5 | 3 | 0.65099 | 0.12496 | 0.0028219 | 0.23973 | 1.0622 |
| 5 | 4 | 0.98009 | 0.15095 | 0.00051565 | 0.48331 | 1.4769 |

==============================================================

C) Percentage of trials with a response

| | SumSq | DF | MeanSq | F | pValue | pValueGG | pValueHF | pValueLB |
|---|---|---|---|---|---|---|---|---|
| (Intercept):AllStimuli | 5089.7 | 4 | 1272.4 | 9.0443 | 2.6656e-05 | 0.00063098 | 0.00011026 | 0.013176 |
| Birdno:AllStimuli | 437.45 | 4 | 109.36 | 0.77735 | 0.54652 | 0.49434 | 0.52595 | 0.39865 |
| Error(AllStimuli) | 5627.4 | 40 | 140.69 | | | | | |

**<u>Posthoc test</u>**

| AllStimuli_1 | AllStimuli_2 | Difference | StdErr | pValue | Lower | Upper |
|---|---|---|---|---|---|---|
| 1 | 2 | 12.389 | 3.6766 | 0.044248 | 0.28937 | 24.489 |
| 1 | 3 | 7.5317 | 3.9818 | 0.37987 | -5.5728 | 20.636 |

| | | | | | | |
|---|---|---|---|---|---|---|
| 1 | 4 | 22.806 | 4.0169 | 0.0014833 | 9.586 | 36.026 |
| 1 | 5 | 48.8 | 6.4393 | 0.00014245 | 27.608 | 69.993 |
| 2 | 1 | -12.389 | 3.6766 | 0.044248 | -24.489 | -0.28937 |
| 2 | 3 | -4.8576 | 3.9027 | 0.72772 | -17.702 | 7.9866 |
| 2 | 4 | 10.417 | 3.1809 | 0.051279 | -0.05177 | 20.885 |
| 2 | 5 | 36.411 | 6.6943 | 0.0020513 | 14.38 | 58.442 |
| 3 | 1 | -7.5317 | 3.9818 | 0.37987 | -20.636 | 5.5728 |
| 3 | 2 | 4.8576 | 3.9027 | 0.72772 | -7.9866 | 17.702 |
| 3 | 4 | 15.274 | 3.5436 | 0.010437 | 3.6121 | 26.937 |
| 3 | 5 | 41.269 | 6.1355 | 0.00038674 | 21.076 | 61.461 |
| 4 | 1 | -22.806 | 4.0169 | 0.0014833 | -36.026 | -9.586 |
| 4 | 2 | -10.417 | 3.1809 | 0.051279 | -20.885 | 0.05177 |
| 4 | 3 | -15.274 | 3.5436 | 0.010437 | -26.937 | -3.6121 |
| 4 | 5 | 25.994 | 5.2096 | 0.0038521 | 8.849 | 43.139 |
| 5 | 1 | -48.8 | 6.4393 | 0.00014245 | -69.993 | -27.608 |
| 5 | 2 | -36.411 | 6.6943 | 0.0020513 | -58.442 | -14.38 |
| 5 | 3 | -41.269 | 6.1355 | 0.00038674 | -61.461 | -21.076 |
| 5 | 4 | -25.994 | 5.2096 | 0.0038521 | -43.139 | -8.849 |

```
===========================================================================
```

**Statistics for Fig. 2**

Results of linear mixed effects model

Fig. 2B - mean number of calls

```
              Mixed Linear Model Regression Results
=================================================================
Model:                 MixedLM    Dependent Variable:   CallResponse
No. Observations:      30         Method:               REML
No. Groups:            3          Scale:                0.0432
Min. group size:       10         Log-Likelihood:       -3.6154
Max. group size:       10         Converged:            Yes
Mean group size:       10.0
-----------------------------------------------------------------
                       Coef.  Std.Err.   z    P>|z| [0.025 0.975]
-----------------------------------------------------------------
Intercept                 0.210    0.132  1.593 0.111 -0.048  0.469
```

```
SongQuality[T.Normal]    0.340    0.085  4.009 0.000  0.174  0.507
INQuality[T.Degraded]   -0.052    0.112 -0.464 0.643 -0.272  0.168
INQuality[T.Normal]     -0.062    0.085 -0.736 0.462 -0.229  0.104
BackgroundNoise[T.Yes]   0.100    0.076  1.317 0.188 -0.049  0.249
Group Var                0.032    0.181
=================================================================
```

Fig. 2B - response latency

```
              Mixed Linear Model Regression Results
=================================================================
Model:                  MixedLM    Dependent Variable:    Latency
No. Observations:       30         Method:                REML
No. Groups:             3          Scale:                 0.1818
Min. group size:        10         Log-Likelihood:        -20.3306
Max. group size:        10         Converged:             Yes
Mean group size:        10.0
-----------------------------------------------------------------
                        Coef.  Std.Err.   z    P>|z| [0.025 0.975]
-----------------------------------------------------------------
Intercept                1.143    0.193  5.909 0.000  0.764  1.522
SongQuality[T.Normal]   -0.429    0.174 -2.462 0.014 -0.770 -0.087
INQuality[T.Degraded]   -0.082    0.230 -0.357 0.721 -0.533  0.369
INQuality[T.Normal]     -0.235    0.174 -1.352 0.176 -0.576  0.106
BackgroundNoise[T.Yes] -0.047    0.156 -0.300 0.764 -0.352  0.258
Group Var                0.026    0.108
=================================================================
```

Fig. 2D - percentage response

```
               Mixed Linear Model Regression Results
===================================================================
Model:               MixedLM  Dependent Variable:  PercentageResponse
No. Observations:    30       Method:              REML
No. Groups:          3        Scale:               221.6153
Min. group size:     10       Log-Likelihood:      -110.5873
Max. group size:     10       Converged:           Yes
Mean group size:     10.0
-------------------------------------------------------------------
                         Coef.  Std.Err.   z    P>|z|  [0.025 0.975]
-------------------------------------------------------------------
Intercept                18.611   10.135  1.836 0.066  -1.253 38.475
SongQuality[T.Normal]    23.611    6.077  3.885 0.000  11.699 35.523
```

```
INQuality[T.Degraded]   -6.944    8.040 -0.864 0.388 -22.702  8.813
INQuality[T.Normal]     -6.944    6.077 -1.143 0.253 -18.856  4.967
BackgroundNoise[T.Yes]   7.222    5.436  1.329 0.184  -3.432 17.876
Group Var              202.874   15.763
==================================================================
```

**<u>Statistics for Figure S1 (Familiar Song Playback Assay)</u>**

Number of birds = 4; Number of stimuli = 5

Stimuli order: 1- IN motif; 2- Only IN; 3- Motif IN; 4- Only motif; 5- Noise

Test used: Repeated measures ANOVA, posthoc: Tukey Karmer

A) Normalised mean number of calls

| | SumSq | DF | MeanSq | F | pValue | pValueGG | pValueHF | pValueLB |
|---|---|---|---|---|---|---|---|---|
| (Intercept):AllStimuli | 0.60521 | 4 | 0.1513 | 2.9686 | 0.088997 | 0.19711 | 0.096214 | 0.22704 |
| Birdno:AllStimuli | 0.16522 | 4 | 0.041305 | 0.81041 | 0.5523 | 0.48409 | 0.54744 | 0.46301 |
| Error(AllStimuli) | 0.40775 | 8 | 0.050968 | | | | | |

==============================================================

==============================================================

B) Mean latency

Calls produced only during stimulus are considered. In case birds didn't respond mean latency 1.77 sec was used for statistics, since that was the maximum possible latency

SumSq DF MeanSq F pValue pValueGG pValueHF pValueLB

| | | | | | | | | |
|---|---|---|---|---|---|---|---|---|
| (Intercept):AllStimuli | 0.18525 | 4 | 0.046313 | 1.0612 | 0.43485 | 0.41176 | 0.41332 | 0.41122 |
| Birdno:AllStimuli | 0.058301 | 4 | 0.014575 | 0.33398 | 0.84779 | 0.6248 | 0.63399 | 0.62172 |
| Error(AllStimuli) | 0.34913 | 8 | 0.043642 | | | | | |

==================================================================================================================================

C) Percentage response

| | SumSq | DF | MeanSq | F | pValue | pValueGG | pValueHF | pValueLB |
|---|---|---|---|---|---|---|---|---|
| (Intercept):AllStimuli | 669.14 | 4 | 167.29 | 0.98476 | 0.46765 | 0.42923 | 0.43913 | 0.42561 |
| Birdno:AllStimuli | 244.56 | 4 | 61.141 | 0.35992 | 0.83056 | 0.62443 | 0.6686 | 0.60947 |
| Error(AllStimuli) | 1359 | 8 | 169.88 | | | | | |
